# Continuous hypermutation and evolution of noncanonical amino acid synthases

**DOI:** 10.64898/2026.03.11.711232

**Authors:** Yuichi Furuhata, Gordon Rix, Patrick J. Almhjell, Chang C. Liu

## Abstract

Genetic code expansion (GCE) enables the site-specific incorporation of noncanonical amino acids (ncAAs) into proteins but is constrained by reliance on exogenously supplied chiral ncAAs. Achieving intracellular ncAA biosynthesis would enable more scalable and cost-effective GCE. Here, we report the continuous hypermutation and evolution of amino acid synthases that produce high levels of ncAAs inside yeast, thus supporting GCE from simple ncAA precursors. We encoded an engineered ‘tyrosine synthase’ (*Tm*TyrS) on an error-prone orthogonal DNA replication system (OrthoRep) and selected variants based on ncAA biosynthesis from readily available phenol analogs and intracellular L-serine. Our selection employed orthogonal ncAA-specific aminoacyl-tRNA synthetases (aaRSs) as biosensors whereby target ncAA production leads to aminoacylation of an amber suppressor tRNA and the translation of a selectable reporter containing an amber stop codon. Our evolution successfully yielded *Tm*TyrS variants that efficiently produced 3-iodo-, 3-bromo-, 3-chloro-, and 3-methyl-L-tyrosine, enabling amber codon-specified ncAA-dependent translation, in some cases at levels comparable to sense codon-specified natural amino acid translation. This work reduces barriers for expressing proteins containing substituted tyrosines. Moreover, because aaRSs can themselves be evolved (including with OrthoRep) for a flexible range of ncAA specificities, these results establish an end-to-end framework for evolving ncAA biosynthetic enzymes *in vivo*.

**Graphical abstract:** 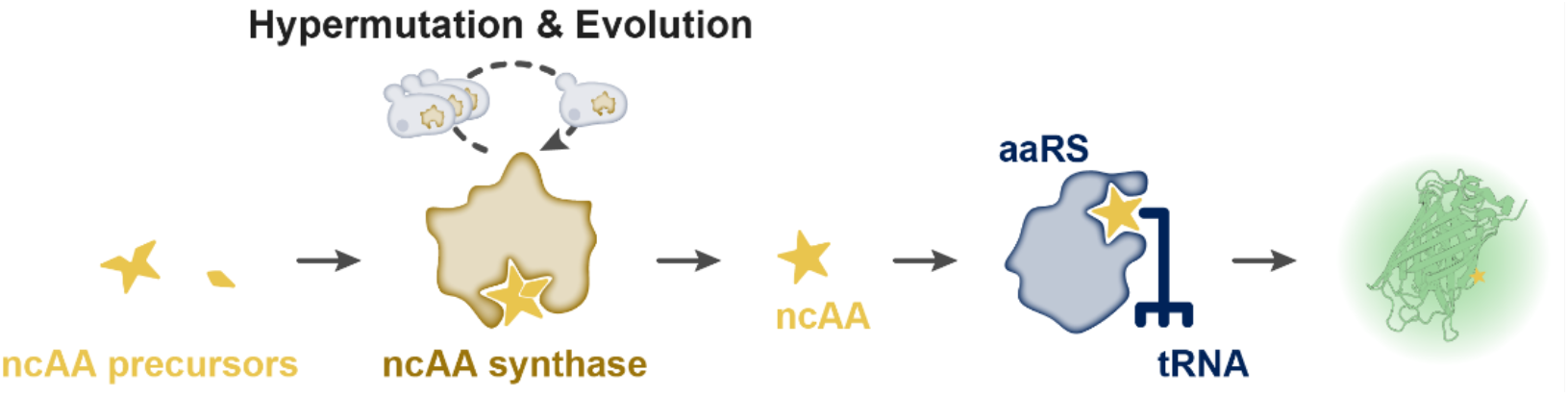

We describe an OrthoRep-driven platform for evolving noncanonical amino acid (ncAA) synthases. Hypermutation of ncAA synthase genes enables evolution of ncAA biosynthesis from simple precursors, while intracellular ncAA production is linked to fluorescence via an orthogonal aaRS/tRNA system, allowing FACS enrichment of improved variants through iterative cycles.

## Introduction

The natural translation machinery uses only 20 canonical amino acids for protein synthesis. Over the past decades, genetic code expansion (GCE) has emerged as a powerful strategy to overcome this rule by enabling the site-specific incorporation of noncanonical amino acids (ncAAs) into proteins in living cells^[1]^. Much of GCE relies on engineered orthogonal aminoacyl-tRNA synthetases (aaRSs) that recognize ncAAs of interest and charge them onto cognate tRNAs that decode blank codons, most commonly amber suppressor tRNAs^[2,3]^. Such engineered aaRS/tRNA pairs have enabled the site-specific translational incorporation of over 200 types of ncAAs^[4–6]^, advancing protein biology^[7–9]^, cell biology^[10,11]^, enzyme engineering^[12–14]^, chemical biology^[15–17]^, imaging^[18–20]^, and therapeutics^[21–23]^.

While engineered/evolved aaRSs have substantially expanded the translational repertoire, a practical bottleneck in many GCE applications is the reliance on externally synthesized ncAAs^[24,25]^. Because ncAAs are usually absent from endogenous cellular metabolism, they must be chemically synthesized and exogenously supplied at high concentrations, imposing constraints on cost, cellular uptake, and scalability. Biosynthesis of ncAAs integrated into cellular metabolism represents a promising strategy to alleviate these bottlenecks by enabling cost-effective and scalable ncAA production that directly feeds GCE systems *in vivo*. To date, a variety of ncAAs have been biosynthesized using native or engineered enzymes, including phenylalanine^[26–28]^, tyrosine^[29,30]^, tryptophan^[31–33]^, and cysteine^[34,35]^ analogs. However, due to the lack of known biosynthetic pathways for most ncAAs and generally low production efficiencies, the diversity of ncAAs accessible via biosynthesis remains significantly limited compared to the scope of engineered/evolved aaRSs^[25]^. Accordingly, establishing a broadly applicable platform to develop ncAA biosynthetic pathways that both expand the accessible ncAA repertoire and improve biosynthetic efficiency remains a major challenge.

We envisioned an end-to-end experimental evolution framework for the creation of organisms with GCEs that biosynthesize their own ncAAs (Fig. 1). In the first part of this framework, aaRSs are evolved to incorporate ncAAs, achieving GCE from exogenously supplied ncAAs. This has been done previously via a number of strategies^[3,36]^ including using orthogonal DNA replication (OrthoRep) to drive continuous aaRS hypermutation, which yielded highly efficient aaRS/tRNA systems for GCE in a streamlined manner^[37]^. In the second part of this framework, evolved aaRSs are repurposed as biosensors that couple *in vivo* ncAA production to the translation of selectable reporters, while amino acid synthases—rather than aaRSs—are encoded on OrthoRep to drive evolution of ncAA biosynthesis from simple precursors. Here, we describe the second part of this framework to complete our envisioned platform by evolving a synthase for the *in vivo* production of noncanonical L-tyrosines (ncTyrs) from readily available phenol analogs. Since ncTyrs constitute a major class of ncAAs used in GCE, this represents an ideal test and demonstration of our framework. We present the evolution of tyrosine synthase^[38]^ (TyrS) variants that efficiently convert simple phenol analogs, including 2-iodo-, 2-chloro-, 2-bromo-, and 2-methylphenol, into their corresponding ncTyrs *in vivo*, which are then directly used in translation of ncAA-containing proteins.

**Figure 1.**
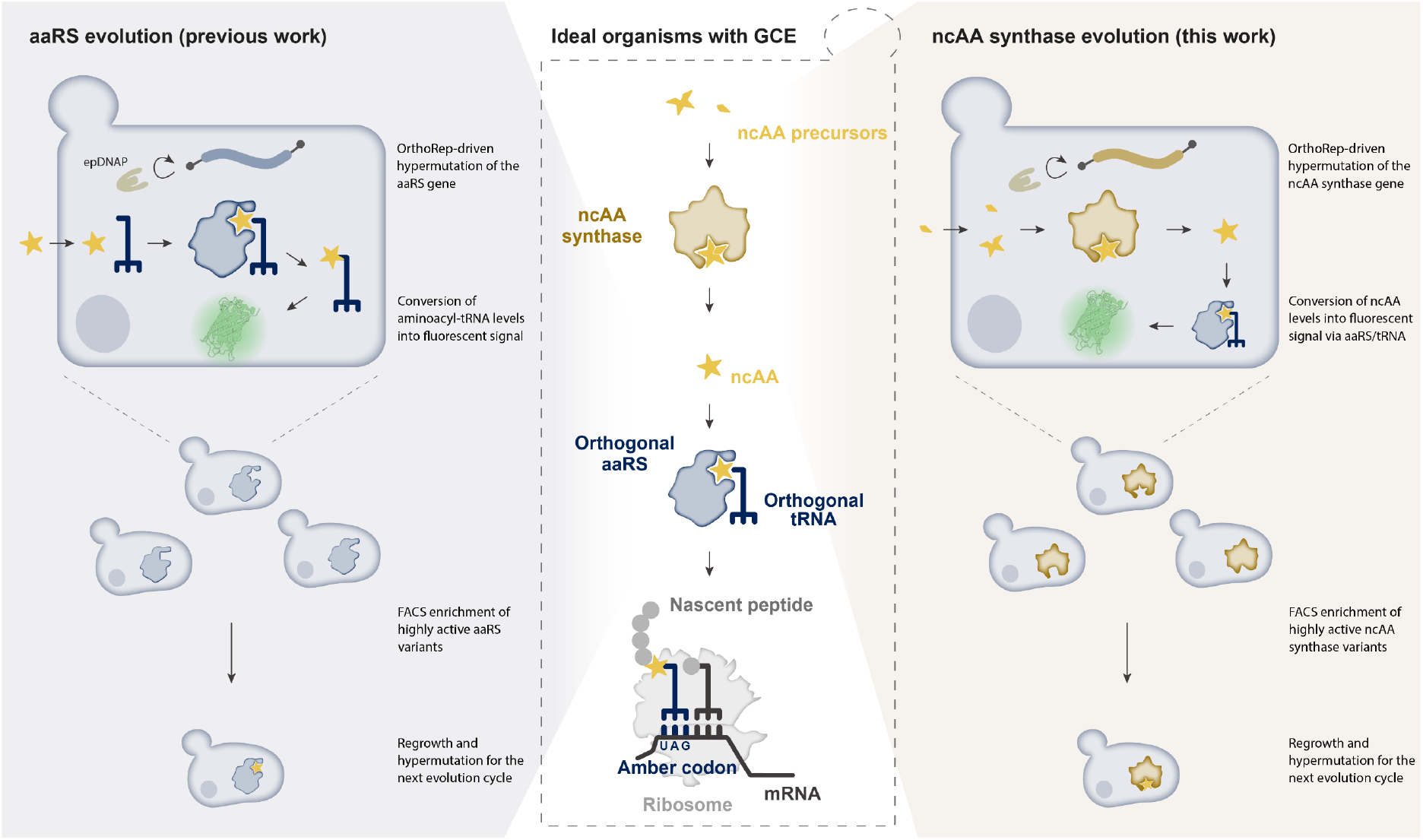
End-to-end experimental evolution framework for the creation of organisms with GCEs that biosynthesize their own ncAAs. Center, conceptual illustration of an ideal GCE-enabled organism. In this organism, ncAA precursors are converted into target ncAAs *in vivo* by an intracellular ncAA synthase. The synthesized ncAA is recognized by an orthogonal aaRS, charged onto its cognate tRNA, and incorporated at a reassigned codon during translation. To establish this architecture, we develop an end-to-end experimental evolution framework that enables evolution of both an orthogonal aaRS and an ncAA synthase within the same platform and workflow. Left, OrthoRep-driven aaRS evolution platform described previously^[37]^. Hypermutation is applied to an orthogonal aaRS on an OrthoRep plasmid to evolve its specificity toward a target ncAA. Intracellular aminoacyl-tRNA levels are coupled to a fluorescent readout, enabling FACS enrichment of improved aaRS variants through iterative cycles. Right, OrthoRep-driven ncAA synthase evolution platform developed in this study. Hypermutation is applied to an ncAA synthase on an OrthoRep plasmid to evolve biosynthetic activity of target ncAAs. Intracellular ncAA production is coupled to a fluorescent readout via orthogonal aaRS/tRNA system, enabling FACS enrichment of improved ncAA synthase variants through iterative cycles.

## Results and Discussions

### Evolution of *Tm*TyrS with OrthoRep

Tyrosine synthase (*Tm*TyrS), an enzyme engineered from the *Thermotoga maritima* tryptophan synthase β-subunit (*Tm*TrpB), catalyzes the formation of diverse ncTyrs from the corresponding phenol analog and L-serine *in vitro*^[38]^. Evolved under high concentrations of substrates, the most active variant identified in a previous study (*Tm*TyrS6) has only moderate catalytic efficiency^[38]^, leaving room for evolutionary improvement in our platform under physiologically relevant substrate concentrations. Our starting point for evolution was *Tm*TyrSc, an *in vivo*-adapted *Tm*TyrS variant evolved from *Tm*TyrS6 through an extensive but incomplete directed evolution campaign for ncTyr biosynthesis in yeast (Supplementary Results and Supplementary Fig. 1). To couple the production of ncTyrs to cellular fitness, we employed NitroY-F5, a *Methanomethylophilus alvus* pyrrolysyl-tRNA synthetase variant that accepts a range of ncTyrs^[39]^, or its OrthoRep-evolved variant NitroY-F5/3FY-D^[37]^ as the biosensor. *Tm*TyrS6 itself exhibited insufficient ncTyr biosynthesis activity from the corresponding substituted phenols to support survival through suppression of an amber stop codon-containing selection marker by tRNAs aminoacylated with ncTyrs by NitroY-F5 (Supplementary Results and Supplementary Fig. 1).

For selection, we adopted our previously reported fluorescence-activated cell sorting (FACS)-based aaRS evolution system based on a ratiometric ‘RXG’ reporter in which RFP and GFP are connected by a linker containing an amber stop codon^[37,40,41]^. In this system, production of ncTyrs increases the amount of aminoacylated tRNA molecules which in turn increases the probability that the amber codon between RFP and GFP is suppressed. This reporter configuration enabled enrichment of cells that sampled highly active *Tm*TyrS variants through mutation by gating on cells exhibiting higher GFP expression relative to RFP in the presence of chosen phenol analogs during sorting. To normalize RXG reporter fluorescence, we measured GFP and RFP fluorescence of cells expressing an ‘RYG’ reporter, in which the amber codon was replaced with a tyrosine-encoding sense codon. Relative readthrough efficiency (RRE) is defined as the ratio of 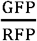 for RXG to that for RYG^[37,40,41]^, such that an RRE of 1 indicates amber suppression is as efficient as native translation at that position. Accordingly, we aimed to evolve *Tm*TyrS variants that exhibited high RREs in the presence of phenol analogs while maintaining low RREs in their absence. Because this selection system directly selects cells based on the readthrough ratio, which must depend on the level of aminoacylated orthogonal tRNA rather than on reporter protein expression levels, it is well suited to select cells with enhanced ncAA production.

To establish the strain for OrthoRep-mediated evolution of *Tm*TyrS, we started with *Saccharomyces cerevisiae* GR-Y567, which carries deletions in *LEU2* and *HIS3*, a split-*LEU2* landing pad p1^[42]^, and a wild-type orthogonal DNA polymerase (DNAP) at the *CAN1* locus^[42]^. We transformed this base strain with the reporter plasmid, which encodes the RXG reporter, an engineered/evolved aaRS, and its orthogonal amber-suppressor tRNA. We then integrated the *Tm*TyrS gene onto the split-*LEU2* landing pad p1 and subsequently replaced the wild-type orthogonal DNAP cassette with error-prone DNAPs (epDNAPs; Fig. 2a) to begin evolution. In the resulting strains, the epDNAP continuously replicates the *Tm*TyrS sequence at a high mutation rate of 1.6 × 10^−5^ and 3.9 × 10^−5^ substitutions per base for BB-Tv and Trixy epDNAPs^[42]^, respectively, while insulating the nuclear genome from the elevated mutagenesis.

**Figure 2.**
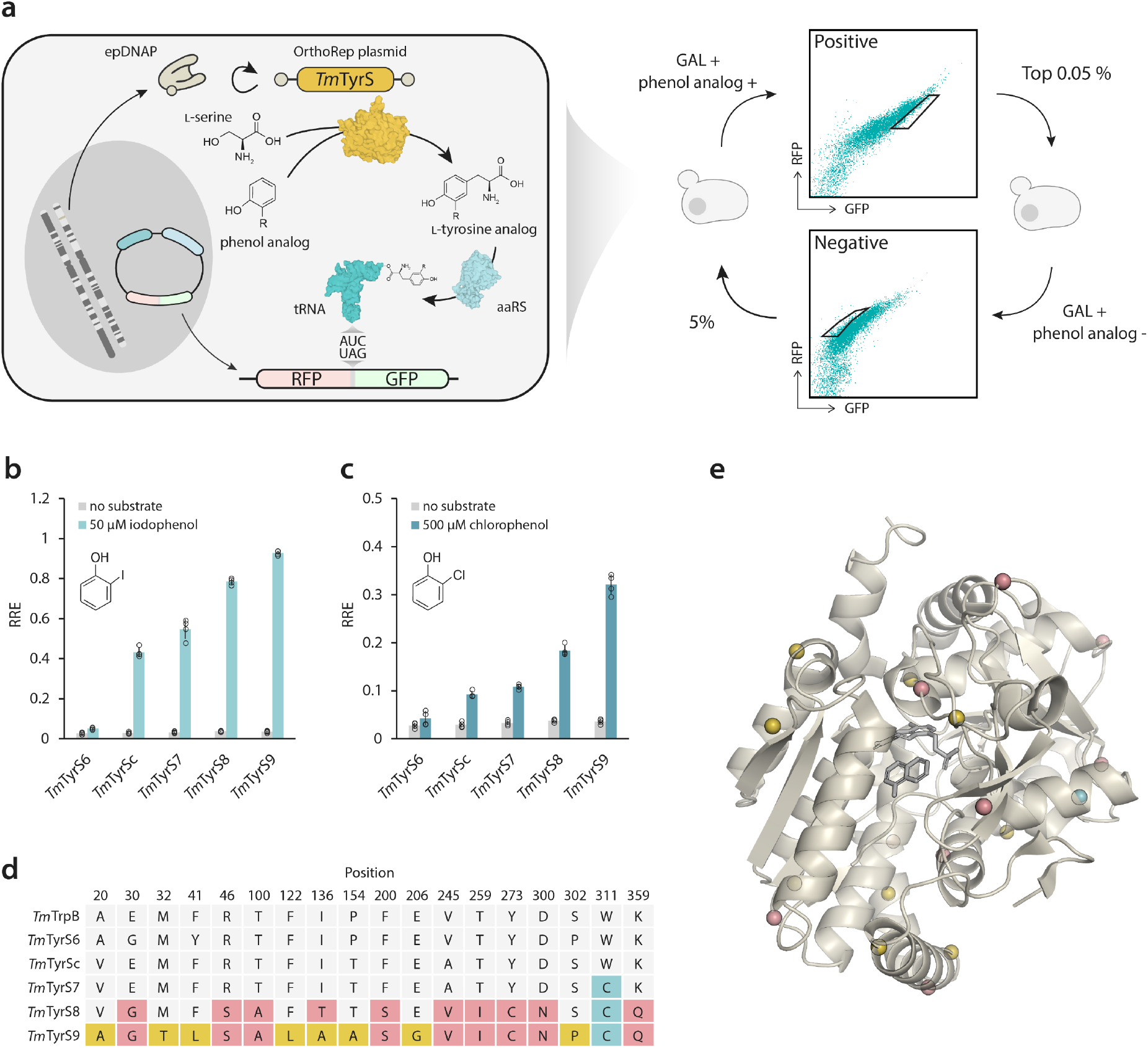
Directed evolution of *Tm*TyrS with OrthoRep. (a) Schematic for a directed evolution approach to evolve the activity of *Tm*TyrS for the synthesis of tyrosine analogs from phenol analogs. (b–c) RRE measurements of the evolved *Tm*TyrS lineage with 2-iodophenol (b) and 2-chlorophenol (c). Each condition was measured in biological quadruplicates, and the mean ± one standard deviation (error bars) is shown. GR-Y261 and pYF227 were used as the base strain and reporter plasmid, respectively. (d) Mutations accumulated in *Tm*TyrS during evolution campaigns. Detailed sequence alignments are provided in Supplementary Data 5. (e) Crystal structure of *Tm*TyrS1 (PDB 8EH1) with mutated residues relative to the parental sequence in the evolution experiments that yielded *Tm*TyrS7 (teal), S8 (salmon pink), and S9 (gold) are highlighted. The colors correspond to those shown in panel (d). *Tm*TyrS1 is the ancestral sequence for all *Tm*TyrS variants described in this study.

Next, we performed iterative FACS-based evolution cycles. After installation of the epDNAP, cells were first grown for approximately 35 generations at 30 °C to allow diversification before starting selection. Each evolution cycle proceeded as follows: (1) induction of the RXG reporter with galactose in the presence or absence of phenol analogs for 48 h at 30 °C, corresponding to ∼3 generations due to slow growth under induction; (2a) a positive selection sort in which ∼10,000,000 cells were screened and the top 0.05% exhibiting the highest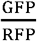 ratio in the presence of phenol analogs were collected, or (2b) a negative selection sort in which ∼1,000,000 cells were screened and the 5% displaying the lowest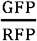 ratio in the absence of phenol analogs were collected; and (3) regrowth of the sorted populations to saturation at 30 °C, corresponding to ∼13–15 generations depending on the number of recovered cells (Fig. 2a). Negative selection was applied every two to four rounds to eliminate cells capable of expressing GFP in the absence of phenol analogs, such as those harboring mutations that converted the amber codon in the RXG reporter to a sense codon. We used both NitroY-F5 and its OrthoRep-evolved variant NitroY-F5/3FY-D as biosensors for ncTyrs. NitroY-F5/3FY-D was evolved with 3-fluoro-L-tyrosine and exhibited higher activity toward many ncTyrs compared to NitroY-F5^[37]^.

As *Tm*TyrS genes on the p1 plasmid autonomously diversified during cultivation, repeated cycles progressively enriched *Tm*TyrS variants with improved activity toward ncTyr biosynthesis. After completing the final cycle, *Tm*TyrS genes were amplified from p1 by PCR, cloned under the TDH3 promoter on a *CEN/ARS* plasmid in a library format, and transformed into a fresh yeast strain carrying the same reporter plasmid. The resulting library underwent a final round of positive selection without hypermutation to observe clonal *Tm*TyrS mutant fitnesses. Approximately 40 clones were then randomly picked and evaluated, and the 4–6 clones exhibiting the highest RRE values with their target phenol analogs (and minimal RRE in their absence) were selected for further characterization and engineering.

Starting from *Tm*TyrSc, we sequentially performed three evolution campaigns and obtained a series of evolved variants (*Tm*TyrS7–9). We chose 2-iodophenol and 2-chlorophenol as evolution substrates, because *Tm*TyrS exhibits catalytic efficiency toward these substrates^[38,43]^ and both NitroY-F5 and NitroY-F5/3FY-D efficiently aminoacylate the orthogonal tRNA with the corresponding reaction products, 3-iodo-L-tyrosine and 3-chloro-L-tyrosine^[37,39]^. The detailed conditions for each evolution experiment and the acquired mutations are provided in Supplementary Data 1 and 2, respectively. *Tm*TyrS8 and *Tm*TyrS9 were generated by combining beneficial mutations identified in parallel evolution experiments with the best-performing variants (Supplementary Fig. 2 and 3). The activities of these *Tm*TyrS variants toward 2-iodophenol and 2-chlorophenol are shown in Figures 2b and 2c. Compared to *Tm*TyrS6, *Tm*TyrSc exhibited enhanced production of 3-iodo-L-tyrosine, which is consistent to the results of the URA3 reporter-based growth assay (Supplementary Information). The production of 3-iodo-L-tyrosine increased progressively as evolution proceeded from *Tm*TyrSc to *Tm*TyrS9 (Fig. 2b). A similar evolutionary trend was observed for 3-chloro-L-tyrosine production, with pronounced improvements in *Tm*TyrS8 and *Tm*TyrS9, which were obtained from evolution experiments using 2-chlorophenol as the substrate (Fig. 2c). Overall, these results demonstrate a substantial enhancement of ncTyr biosynthesis through the OrthoRep-mediated evolution campaigns.

### Evolutionary outcomes

Figures 2d and 2e show the amino acid substitutions accumulated during the FACS-based evolution campaigns. Using *Tm*TyrSc as a reference sequence, mutations acquired sequentially in *Tm*TyrS7 (teal, 1 substitution), *Tm*TyrS8 (salmon pink, 10 substitutions), and *Tm*TyrS9 (gold, 8 substitutions) are highlighted, reflecting their stepwise evolutionary accumulation. In total, the final variant *Tm*TyrS9 harbors 19 cumulative amino acid substitutions through the OrthoRep-mediated FACS-based evolution campaign described here.

Among the 19 new substitutions, D300N is particularly notable. This residue is located within the catalytic pocket of *Tm*TyrS and highly conserved among TrpB enzymes, found as either aspartate (D300), in biosynthetic TrpB enzymes, or arginine (R300), in a class of stand-alone, indole-scavenging TrpB2 enzymes^[44]^. The asparagine (N300) substitution obtained through this OrthoRep-mediated evolution campaign is practically absent from the TrpB evolutionary record, present in only 0.33% (60/18,051) of TrpB-like sequences in a previously compiled dataset^[38]^ (Supplementary Table 1 and Supplementary Data 3). This position is known to be involved in both catalytic and allosteric mechanisms. Its sidechain interacts directly with the hydroxyl group of the L-serine substrate (or, in TrpB2s, the phosphate group of the phospho-L-serine substrate) when it is bound to the pyridoxal 5’-phosphate (PLP) cofactor^[45]^. In TrpBs, its binding to T292 is associated with an allosteric transition induced by its binding partner TrpA, and transitions between binding T292 and L-serine substrate are associated with a closed conformation of the enzyme that generates its reactive aminoacrylate intermediate^[45,46]^. The substitution T292S, present in all engineered *Tm*TyrS variants, is known to recapitulate this allosteric effect in the absence of TrpA, enhancing TrpB’s stand-alone activity^[47]^. While the consequences of converting aspartate’s carboxylate sidechain to a carboxamide—and how this may or may not be specifically activating for tyrosine synthesis—remain unclear, D300N was enriched in two of four independent *Tm*TyrS7 evolution experiments (Supplementary Data 2), suggesting an important role in enhanced TyrS activity.

Also notable is the F200S substitution, as this position is dominated by F (∼70%, in TrpBs) and H (∼20%, in TrpB2s). S200 is present in only 0.27% (48/18,051) of the TrpB-like sequences (Supplementary Table 2 and Supplementary Data 4). F200 interacts with the helix that bears the catalytic lysine, a residue that is used to bind PLP and hypothesized to catalyze a concerted proton transfer during the aminoacrylate-mediated alkylation of phenol by TyrS enzymes.

Since we obtained *Tm*TyrSc by shuffling yeast-adapted *Tm*TrpB sequences with *Tm*TyrS6 (Supplementary Information), it is possible that *Tm*TyrSc lost some beneficial mutations for TyrS activity. Interestingly, of the 19 substitutions acquired during the evolution from *Tm*TyrSc to *Tm*TyrS9, four mutations (V20A, E30G, A245V, and S302P) represent reversions to pre-*Tm*TyrSc residues (Fig. 2d and Supplementary Data 5). V20A and A245V represent reversions to residues present in both wild-type *Tm*TrpB and *Tm*TyrS6. In contrast, E30G and S302P correspond to substitutions identified during previous engineering efforts: E30G was identified during engineering of *Tm*TrpB for reduced-temperature tryptophan synthesis^[48]^, while S302P was acquired during the directed evolution campaign for tyrosine synthase activity^[38]^. These reversion mutations are likely to play important roles in TyrS activity in the evolved variants, whether by retaining important features for aminoacrylate generation used by all TrpBs or for phenol alkylation used by the new TyrS enzymes.

### Characteristics of the evolved *Tm*TyrS

*Tm*TyrS6, the immediate precursor of *Tm*TyrSc, has previously been reported to produce ncTyrs not only from 2-iodophenol and 2-chlorophenol but also from a variety of phenol analogs, including 2-bromophenol and 2-methylphenol^[38,43]^. Because these phenol derivatives are highly similar in structure, we hypothesized that *Tm*TyrS9, which was evolved using 2-iodophenol and 2-chlorophenol, might also exhibit high activity toward other phenol analogs. To test this, we compared the activities of *Tm*TyrS9 and *Tm*TyrS6 toward various phenol analogs.

To reconfirm each ncTyr could be detected by the reporter system, commercially available ncTyrs corresponding to the reaction products from each phenol analog were supplied to reporter cells. In all cases, ncTyr-dependent reporter responses were observed (Supplementary Fig. 4). Next, yeast strains harboring *Tm*TyrS9 and the reporter system were evaluated in the presence of phenol analogs at concentrations up to 500 μM, and RRE values were determined across all concentrations. The same experiments were performed for the parental *Tm*TyrS6 for comparison. *Tm*TyrS9 surpassed *Tm*TyrS6 toward all tested substrates including non-evolution substrates 2-bromophenol and 2-methylphenol (Fig. 3a–d). Notably, the addition of 50 μM 2-iodophenol resulted in an RRE of ∼1, which is comparable to that obtained by direct supplementation of 50 μM 3-iodo-L-tyrosine (Fig. 3a). *Tm*TyrS9 also showed high activity toward 2-bromophenol, achieving an RRE of ∼1 with 158 μM bromophenol, which is comparable to that obtained with 50 μM 3-bromo-L-tyrosine (Fig. 3b). The activities of *Tm*TyrS6 toward 2-chlorophenol and 2-methylphenol were nearly undetectable, while *Tm*TyrS9 showed clear responses, with RRE values of 0.36 and 0.23, at 500 μM 2-chlorophenol and 2-methylphenol, respectively (Fig. 3c and d). Given that 3-methyl-L-tyrosine is chemically difficult to synthesize^[43,49,50]^ and thus highly costly (∼1,600 USD g^−1^), while its precursor 2-methylphenol is inexpensive (<0.1 USD g^−1^), *Tm*TyrS9-mediated biosynthesis of 3-methyl-L-tyrosine offers substantial cost reductions. Moreover, *Tm*TyrS catalyzes ncTyr biosynthesis in an effectively irreversible manner by avoiding the thermodynamically favorable degradation of ncTyrs to phenols, pyruvate, and ammonia^[38]^, thereby facilitating efficient coupling of ncTyr production with GCE.

**Figure 3.**
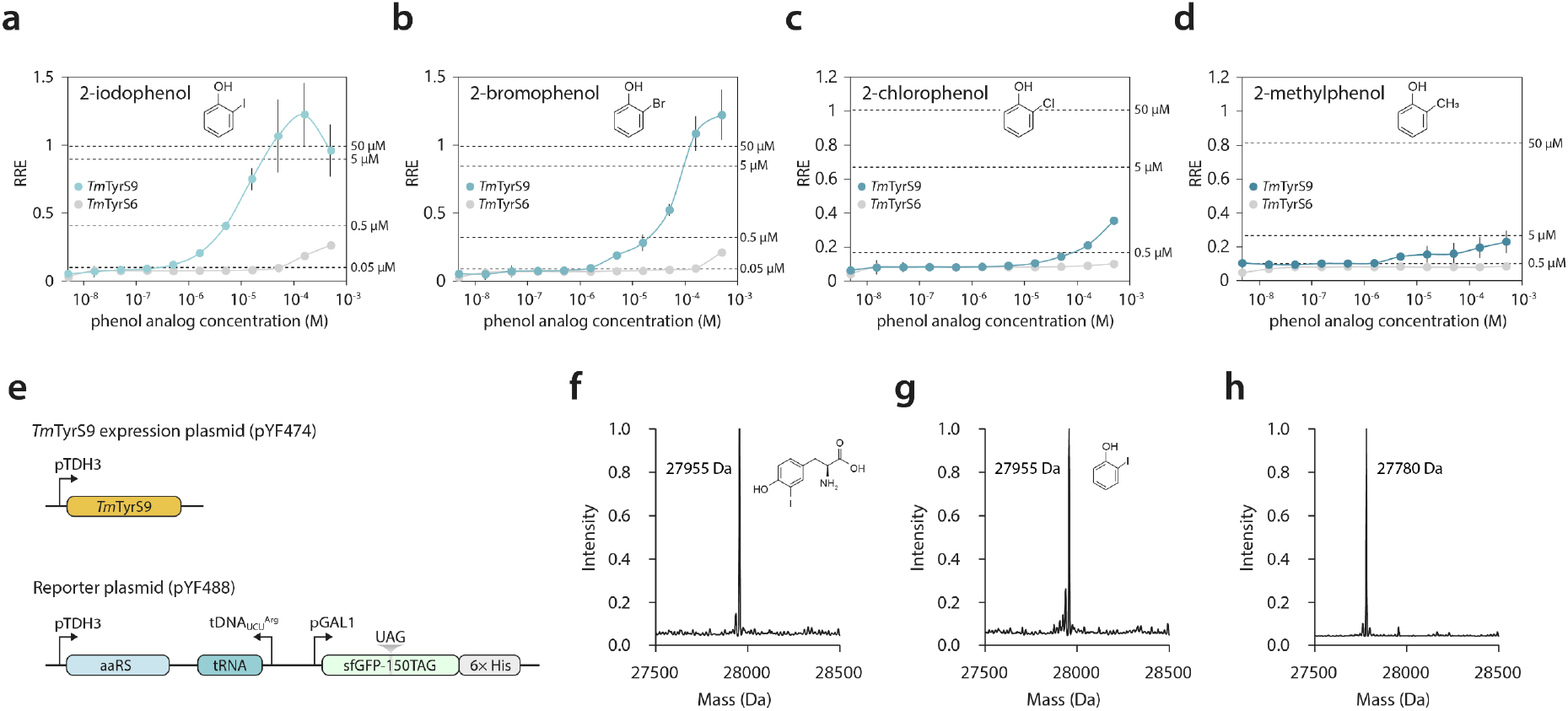
Characterization of *Tm*TyrS9. (a–d) Sensitivity to phenol concentration of *Tm*TyrS6 (gray) and *Tm*TyrS9 (colored). 2-iodophenol (a), 2-bromophenol (b), 2-chlorophenol (c), and 2-methylphenol were titrated. A dotted line in each graph represents the RRE value obtained when the indicated concentration of the corresponding tyrosine analog was added instead of the phenol analog. Each condition was measured in biological quadruplicates, and the mean ± one standard deviation (error bars) is shown. GR-Y261 and pYF227 were used as the base strain and reporter plasmid, respectively. Schematic of the plasmids used for sfGFP150TAG protein expression and purification for protein mass spectrometry. (f–h) Whole-protein mass spectrometry of sfGFP150TAG (f, g) and WT sfGFP (h). sfGFP150TAG was expressed with *Tm*TyrS9 with 500 μM 3-iodo-L-tyrosine (f) or 500 μM 2-iodophenol (g). WT sfGFP was expressed with the empty vector. GR-Y261 and pYF488 were used as the base strain and reporter plasmid, respectively. Source data are provided in Supplementary Data 9.

To evaluate the fidelity of ncTyr biosynthesis mediated by *Tm*TyrS9, we expressed sfGFP containing an amber codon at position 150 together with *Tm*TyrS9 and NitroY-F5/3FY-D in the presence of either 2-iodophenol or 3-iodo-L-tyrosine (Fig. 3e–h). The resulting sfGFP proteins were purified and analyzed by whole protein mass spectrometry. The sfGFP produced in the presence of 3-iodo-L-tyrosine exhibited the expected molecular mass corresponding to incorporation of 3-iodo-L-tyrosine at position N150 (Fig. 3f). Importantly, the sfGFP purified from cultures supplied with 2-iodophenol displayed an identical molecular mass, demonstrating that *Tm*TyrS9 indeed produced 3-iodo-L-tyrosine *in vivo* with high fidelity (Fig. 3g).

## Conclusion

In this work, we extended an OrthoRep-driven aaRS evolution platform to enable continuous hypermutation and evolution of ncAA synthases by repurposing aaRSs as biosensors for ncAA production. Using this platform, we demonstrated the evolution of *Tm*TyrS for the biosynthesis of ncTyrs, resulting in several outcomes including *Tm*TyrS9, which supports the efficient production of ncTyrs from 2-iodophenol, 2-chlorophenol, 2-bromophenol, and 2-methylphenol. Recently, other work that exploits aaRS-based biosensing to evolve ncAA synthases have been reported. Rubini *et al*. demonstrated the directed evolution of a carbamoylase catalyzing the conversion of *N*-carbamoyl-L-3-nitrotyrosine to 3-nitro-L-tyrosine^[51]^. Similarly, Pulschen and Booth *et al*. reported the evolution of a tryptophan halogenase using phage-assisted continuous evolution (PACE)^[52]^ based on aaRSs that recognize tryptophan analogs. While these approaches are related to ours, our OrthoRep-driven platform is distinct in its coupling of *in vivo* continuous hypermutation with biosynthetic selection in yeast cells and its completion of an end-to-end framework where both the aaRS and the ncAA synthase are evolved using the same OrthoRep-driven strategy, offering a unified platform for obtaining expanded genetic code organisms that biosynthesize their own ncAAs for GCE.

The effectiveness of OrthoRep-driven *Tm*TyrS evolution is underscored by two key results; 1) ncTyr biosynthesis mediated by *Tm*TyrS9 in some cases yielded translation efficiencies, as measured by RRE, comparable to those with sense codons; and 2) *Tm*TyrS9 enabled biosynthesis of 3-methyl-L-tyrosine, a compound whose market price exceeds that of its precursor by more than three orders of magnitude. These results demonstrate the practical utility of the ncAA synthases obtained here in addition to proving a general framework for ncAA synthase and synthetase evolution for the GCE field.

## Methods

### DNA plasmid construction

Plasmids used in this study are listed in Supplementary Data 6 together with their DNA sequences. All DNA templates for PCR were obtained from previous studies or synthesized as gBlocks (IDT). All primers were synthesized by IDT. Amplicons for plasmid construction were generated using KOD One PCR Master Mix -Blue- (Toyobo). Plasmids were assembled using either Gibson Assembly or Golden Gate Assembly and transformed into chemically competent or electrocompetent *E. coli* TOP10 cells (ThermoFisher). All plasmids were sequence verified by Sanger sequencing (Azenta) or whole plasmid sequencing (Plasmidsaurus).

### Reagents

All ncAA stock solutions were prepared at a final concentration of 10 mM of the L-isomer. Deionized water was added to solid ncAA to approximately 90% of the final volume, and the pH was gradually adjusted with NaOH as needed to dissolve the ncAA. Solutions were sterile filtered through 0.2 μm filters and stored at −80 °C. After thawing, stocks were stored at 4 °C for up to 8 weeks prior to use. Commercially available key reagents are listed in Supplementary Data 7.

### Yeast strains and media

All yeast strains used in this study are listed in Supplementary Data 8. Yeast was incubated at 30 °C and typically cultured in synthetic complete (SC) medium (20 g/L dextrose, 6.7 g/L yeast nitrogen base w/o amino acids (US Biological), and the appropriate dropout mix (US Biological)) or in MSG (L-Glutamic acid monosodium salt) based SC medium (20 g/L dextrose, 1.72 g/L yeast nitrogen base w/o ammonium sulfate w/o amino acids (US Biological), appropriate nutrient drop-out mix (US Biological), 1 g/L L-Glutamic acid monosodium salt hydrate (ThermoFisher)). Media lacking specific nutrients are denoted as −X, where X indicates the single letter amino acid code of the omitted amino acid or uracil (U). For GAL1 promoter induction, SCGR medium containing 2% galactose and 2% raffinose in place of glucose was used. For selection of the MET15 marker, cells were propagated in media lacking both methionine and cysteine. Liquid cultures (500 μL) in 96-well deep-well plates were incubated at 750 rpm, while all other liquid cultures were incubated at 200 rpm. Agar plates were prepared by mixing equal volumes of 2× molten agar and 2× medium. Prior to experiments, cells were grown to saturation in selective media to maintain plasmids.

### Yeast transformations

All yeast transformations, including p1 integrations and polymerase replacement integrations, were performed using frozen competent cells as previously described^[53]^. For p1 and polymerase replacement integrations, 1–5 μg of plasmid DNA was linearized using ScaI-HF or EcoRI-HF (NEB), respectively. For CEN/ARS plasmid transformations, 100–500 ng of plasmid DNA was used. Transformants were selected on the appropriate selective agar plates. MSG SC−HL agar plates supplemented with 100 mg/L nourseothricin and 200 mg/L L-canavanine were used for polymerase replacement integration. Plates were incubated at 30 °C for 2 days for nuclear plasmid transformations and genomic integrations, and for 4 days for p1 integrations.

All linearized plasmids for p1 integration were integrated into a split LEU2 landing pad to generate the desired p1 constructs^[42]^. Genomic DNA and p1/p2 plasmids were extracted as previously described^[33]^. Briefly, 1.5 mL of yeast culture was pelleted, washed with 0.9% NaCl, and resuspended in 250 μL Zymolyase solution (0.9 M D-sorbitol, 0.1 M EDTA, 10 U/mL Zymolyase (US Biological)). After incubation at 37 °C for 1 h, cells were lysed with proteinase K solution and treated at 65 °C for 30 min. Following potassium acetate precipitation and ethanol precipitation, nucleic acids were resuspended in TE buffer, treated with RNase A, and reprecipitated with isopropanol. The final pellet was resuspended in 30 μL water. Proper integration was confirmed by agarose gel electrophoresis of recombinant p1 DNA. The presence of recombinant p1 was also confirmed after polymerase replacement and evolution campaigns.

### FACS-based *Tm*TyrS evolution and selection with OrthoRep

Prior to each round of FACS selection, yeast strains harboring *Tm*TyrS on p1, a reporter plasmid, and an error-prone orthogonal DNAP (BB-Tv or Trixy)^[42]^ integrated at the CAN1 locus were grown in SC−HL at 30 °C to saturation. Cultures were diluted to OD_600_ = 0.6 in 2 mL medium and grown to OD_600_ = 1.5– 3 (4–7 h). The cells were then induced in SCGR−HL at OD_600_ = 0.6 supplemented with 20 mM L-serine and with or without phenol analogs. Cultures were incubated at 30 °C for 2 days. After culture saturation, the cells were washed and resuspended in HBSM buffer (20 mM HEPES pH 7.5, 150 mM NaCl, 5 mM maltose) and sorted using a Sony SH800S cell sorter with a 100 μm sorting chip. For positive selection, cells were grown with phenol analogs, whereas for negative selection, no phenol analog was supplied. Approximately 10 million (positive) or 1 million (negative) events were measured per sort. GFP and RFP were detected using 488 nm excitation with 525/50 and 617/30 filters, respectively. The top 0.05% of cells positive for both GFP and RFP were recovered for positive sorts while the top 5% of cells positive for RFP and most negative for GFP were recovered for negative sorts. Cells were recovered in 2 mL of SC−HL at 30 °C until saturation and were used for the next round.

After multiple selection rounds, *Tm*TyrS genes were amplified from p1 and subcloned into *CEN/ARS* plasmids to prevent further hypermutation. Libraries were reintroduced into yeast strains with a reporter plasmid and subjected to final positive FACS selections. Approximately 1 million (positive) events were measured per sort. The top 0.1% of cells positive for both GFP and RFP were recovered for positive sorts. After positive sorts, cells were grown on SC−HL media agar plates to isolate individual clones for sequencing and further characterization.

### Flow cytometry

Cells were grown in SC−HL at 30 °C to saturation. Cultures were diluted to OD_600_ = 0.6 in 500 μL and grown to OD_600_ = 1.5–3 (4–7 h). The cells were then induced in 200 μL of SCGR−HL at OD_600_ = 0.6 supplemented with 20 mM L-serine and with or without phenol analogs. The induced cultures were incubated at 30 °C for 2 days. After culture saturation, cells were diluted into 0.9% NaCl and analyzed on an Attune NxT flow cytometer (Life Technologies). The fluorescence of RFP and GFP from 20,000 single cells was recorded, and the mean fluorescence for each population was determined. Data were analyzed using FlowJo v10.10.0. Autofluorescence of cells was subtracted using uninduced cells grown in SC media. Fold changes and RREs were calculated as previously described^[37]^. Cells transformed with the plasmid encoding the RYG reporter for RRE calculation were induced in the absence of phenol analog.

### Protein purification and mass spectrometry

*Tm*TyrS9-expressing yeast harboring an sfGFP-150TAG reporter was grown in SC−HL media to saturation. The culture was diluted to OD_600_ = 0.6 in 20 mL and grown to OD_600_ = 1.5–3 (4–7 h). The cells were then induced in 40 mL of SCGR−HL media at OD_600_ = 0.6 supplemented with 2-iodophenol or 3-iodo-L-tyrosine. The induced cultures were incubated for 2 days. After culture saturation, cells were washed with 0.9% NaCl. Proteins were extracted from yeast cells using Y-PER (ThermoFisher) containing cOmplete, EDTA-free protease inhibitor cocktail (MilliporeSigma), purified using HisPur Ni-NTA resin (ThermoFisher) and eluted with elution buffer (20 mM sodium phosphate, 300 mM NaCl, 250 mM imidazole, pH 8.0). Intact proteins were analyzed by LC/MS (ACQUITY UPLC H-class system and Xevo G2-XS QTof, Waters). Proteins were separated using an ACQUITY UPLC BEH Phenyl VanGuard Pre-column (130Å, 1.7 μm, 2.1 mm × 5 mm, Waters) at 45 °C. The 5-minute method used 0.2 mL/min flow rate of a gradient of Buffer A consisting of 0.1% formic acid in water and Buffer B, acetonitrile. The Xevo Z-spray source was operated in positive MS resolution mode, 400–4,000 Da with a capillary voltage of 3000 V and a cone voltage of 40 V (NaCsI calibration, Leu-enkephalin lock-mass). Nitrogen was used as the desolvation gas at 350 °C and a total flow of 800 L/h. Total average mass spectra were reconstructed from the charge state ion series using the MaxEnt1 algorithm from MassLynx software (Waters) according to the manufacturer’s instructions. To obtain the ion series described, the major peak of the chromatogram was selected for integration before further analysis. The theoretical molecular weight of a protein with 3-iodo-L-tyrosine was calculated by first computing the theoretical molecular weight of wild-type sfGFP and then manually correcting for the theoretical molecular weight of 3-iodo-L-tyrosine.

### Statistics and reproducibility

Microsoft Excel was used for all statistical analyses. Replicate numbers are provided in the figure legends. No statistical methods were used to predetermine sample size, and no data were excluded.

## Data Availability

All data related to evolution campaigns, mutations, sequence alignments, reagents, plasmids, yeast strains, and mass spectrometry are provided in the Supplementary Data.

## Acknowledgements

This work was funded by NIH R35GM136297 (C.C.L.), a JSPS Overseas Research Fellowship 202260318 (Y.F.), Grant in Aid for Scientific Research (B) 25K00105 (Y.F.), and NIH 1F32GM156066 (P.J.A.)

## Author Contributions

Y.F., G.R., and C.C.L. designed the experiments. Y.F. and G.R. conducted the experiments. Y.F., G.R., P.J.A., and C.C.L. analyzed the data and wrote the manuscript.

