## Supplementary for "Continuous hypermutation and evolution of noncanonical amino acid synthases"

### Supplementary Results

#### Growth-based screening of *Tm*TyrS variants adapted for *in vivo* activity

*Tm*TyrS6, a previously obtained *Tm*TrpB variant through conventional directed evolution, is an enzyme capable of producing multiple noncanonical L-tyrosines (ncTyrs) including 3-halogenated L-tyrosines *in vitro*[1,2]. To evaluate the *in vivo* ncTyr biosynthetic activity of *Tm*TyrS6, we introduced *Tm*TyrS6 into *Saccharomyces cerevisiae* together with NitroY-F5[3], an aminoacyl-tRNA synthetase capable of accepting a broad range of ncTyrs including 3-halogenated L-tyrosines, its cognate tRNA, and a URA3 stop-codon suppression reporter. In this system, ncTyrs produced by *Tm*TyrS are charged onto the orthogonal tRNA, thereby increasing URA3 expression in the presence of phenol analogs and resulting in a growth advantage under uracil-deficient conditions. Exogenous supplementation of 3-chloro-L-tyrosine restored the growth of the strain in media lacking uracil, whereas there was no survival dependent on 2-iodophenol (Figure S1a). This finding suggested that *Tm*TyrS6 may not be optimal for *in vivo* activity in yeast due to its extensive evolution under *in vitro* conditions.

To adapt *Tm*TyrS6 for *in vivo* activity in yeast, we compared its sequence with a *Tm*TrpB library previously domesticated for yeast using OrthoRep[4,5]. Four mutations (E30G, I69V, K96L, and L213P) that were highly conserved during yeast evolution but altered in *Tm*TyrS6 were reverted, and three mutations (A20T, A118V, and N167D) frequently observed in the yeast-adapted library were introduced. The resulting variant was designated as *Tm*TyrS6b. *Tm*TyrS6b was subsequently subjected to *in vitro* homologous recombination with the yeast-adapted *Tm*TrpB library to generate a mutant library. This library was subjected to two rounds of positive selection on uracil-deficient solid medium supplemented with 2-iodophenol, interspersed with one round of negative selection using 5-fluoroorotic acid (5-FOA). From this screening, we identified *Tm*TyrSc, which restores growth in uracil-deficient medium in the presence of 2-iodophenol (Figure S1b). Overall, *Tm*TyrSc contains 28 amino acid substitutions relative to *Tm*TyrS6. Details of the screening procedure are described elsewhere[6].

### Supplementary Methods

#### Plating assays

For plating assays with a larger plate surface area, GR-Y261-based yeast strains harboring *Tm*TyrS variants and a reporter plasmid (pGR642) were grown in SC−HL at 30 °C to saturation. An aliquot (100 µL) of each culture was plated onto the appropriate solid medium in a 10 cm petri dish. After 3 days of growth, plates were imaged (Bio-Rad ChemiDoc). Resulting images were adjusted uniformly.

#### Growth-based *Tm*TyrS screening

GR-Y261-based yeast strains harboring the recombination library and a reporter plasmid (pGR642) were grown in SC−HL at 30 °C to saturation. For positive selections, cells were grown on solid SC−HLU media supplemented with 20 mM L-serine and 500 µM 2-iodophenol. For negative selection, cells were grown on solid SC−HL media supplemented with 1 g/L 5-FOA. After the second positive selection, cells were streaked on SC−HL media agar plates to isolate individual clones for sequencing and further characterization.

### Supplementary Tables

**Supplementary Table 1. Amino acid distribution at the *Tm*TrpB D300-equivalent position across 18,051 TrpB-like sequences**

| Amino acid | Counts | Percent (%) | Note |
|---|---|---|---|
| - | 485 | 2.69 | |
| A | 14 | 0.08 | |
| C | 4 | 0.02 | |
| D | 13491 | 74.74 | TrpBs |
| E | 79 | 0.44 | |
| G | 17 | 0.09 | |
| H | 5 | 0.03 | |
| I | 1 | 0.01 | |
| K | 3 | 0.02 | |
| N | 60 | 0.33 | *Tm*TyrS9 |
| Q | 14 | 0.08 | |
| R | 3840 | 21.27 | TrpB2s |
| S | 3 | 0.02 | |
| V | 30 | 0.17 | |
| X | 5 | 0.03 | |

Amino acid frequencies at the position corresponding to D300 in *Tm*TrpB were determined from an alignment of 18,051 TrpB-like sequences. – indicates sequences not aligned at the position. X indicates an unidentified residue.

**Supplementary Table 2. Amino acid distribution at the *Tm*TrpB F200-equivalent position across 18,051 TrpB-like sequences**

| Amino acid | Counts | Percent (%) | Note |
|---|---|---|---|
| - | 3 | 0.02 | |
| A | 12 | 0.07 | |
| C | 1 | 0.01 | |
| E | 1 | 0.01 | |
| F | 12797 | 70.89 | TrpBs |
| H | 3884 | 21.52 | TrpB2s |
| I | 44 | 0.24 | |
| L | 1030 | 5.71 | |
| M | 39 | 0.22 | |
| N | 1 | 0.01 | |
| P | 10 | 0.06 | |
| Q | 8 | 0.04 | |
| S | 48 | 0.27 | *Tm*TyrS9 |
| T | 5 | 0.03 | |
| V | 1 | 0.01 | |

Amino acid frequencies at the position corresponding to F200 in *Tm*TrpB were determined from an alignment of 18,051 TrpB-like sequences. – indicates sequences not aligned at the position. X indicates an unidentified residue.

### Supplementary Figures

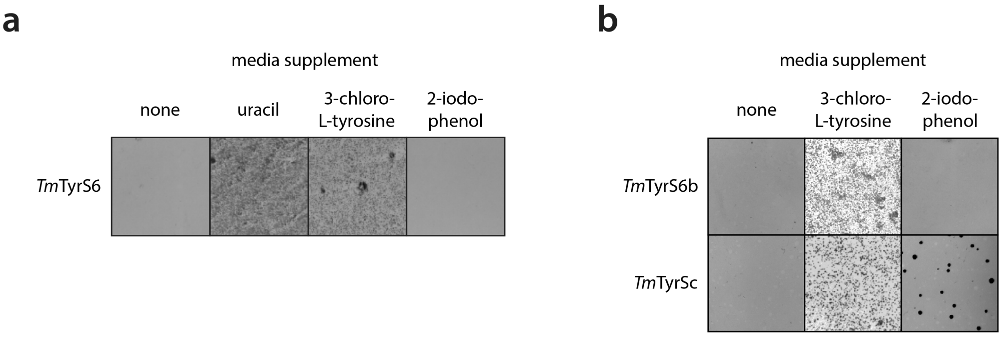


**Supplementary Fig. 1.** Screening of *in vivo*-adapted *Tm*TyrS6 variant. (a) Representative images of yeast expressing *Tm*TyrS6, NitroY-F5, its cognate tRNA, and a URA3 stop-codon suppression reporter grown on SC−HLU agar plates supplemented with no additive, 76 mg/L uracil, 10 mM 3-chloro-L-tyrosine, or 500 µM 2-iodophenol for 3 days. (b) Representative images of yeast expressing the indicated *Tm*TyrS variants together with NitroY-F5, its cognate tRNA, and a URA3 stop-codon suppression reporter grown on SC−HLU agar plates supplemented with no additive, 10 mM 3-chloro-L-tyrosine, or 500 µM 2-iodophenol for 3 days.

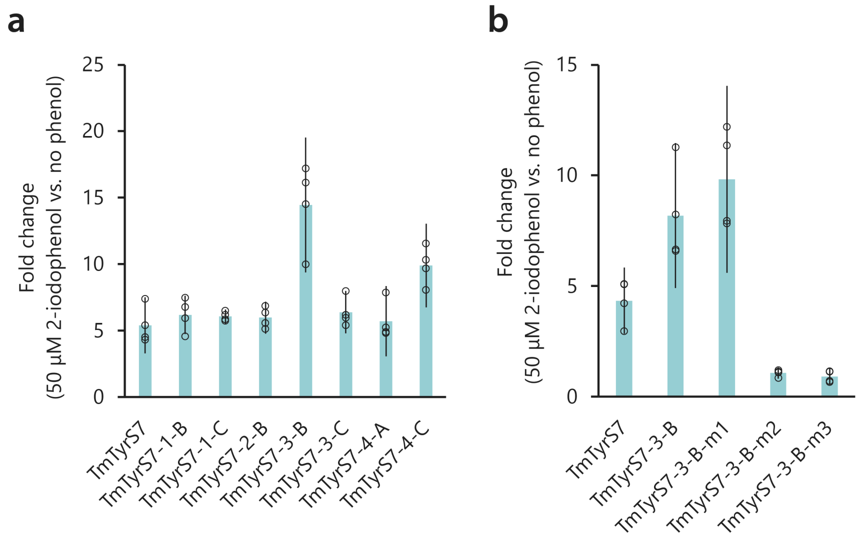


**Supplementary Fig. 2.** Characterization and engineering of *Tm*TyrS7-derived variants in response to 2-iodophenol. (a) Fold increase in RREs relative to no phenol in yeast cells expressing variants obtained from the *Tm*TyrS7 evolution campaign in the presence of 50 μM 2-iodophenol. (b) Fold increase in RREs relative to no phenol for engineered variants based on *Tm*TyrS7-3-B, the best performing variant in panel (a). Beneficial mutations were extracted from other variants that exhibited enhanced responses relative to *Tm*TyrS7, yielding the substitution sets m1 (R46S, T259I, K359Q), m2 (K72E, K143R), and m3 (R46S, K72E, K143R, T259I, K359Q; corresponding to m1 + m2). *Tm*TyrS7-3-B-m1, the best performing variant, was designated as *Tm*TyrS8. RRE values measured in the presence of 50 μM 2-iodophenol were normalized to those measured in its absence. Each condition was measured in biological quadruplicates, and the mean ± one standard deviation (error bars) is shown.

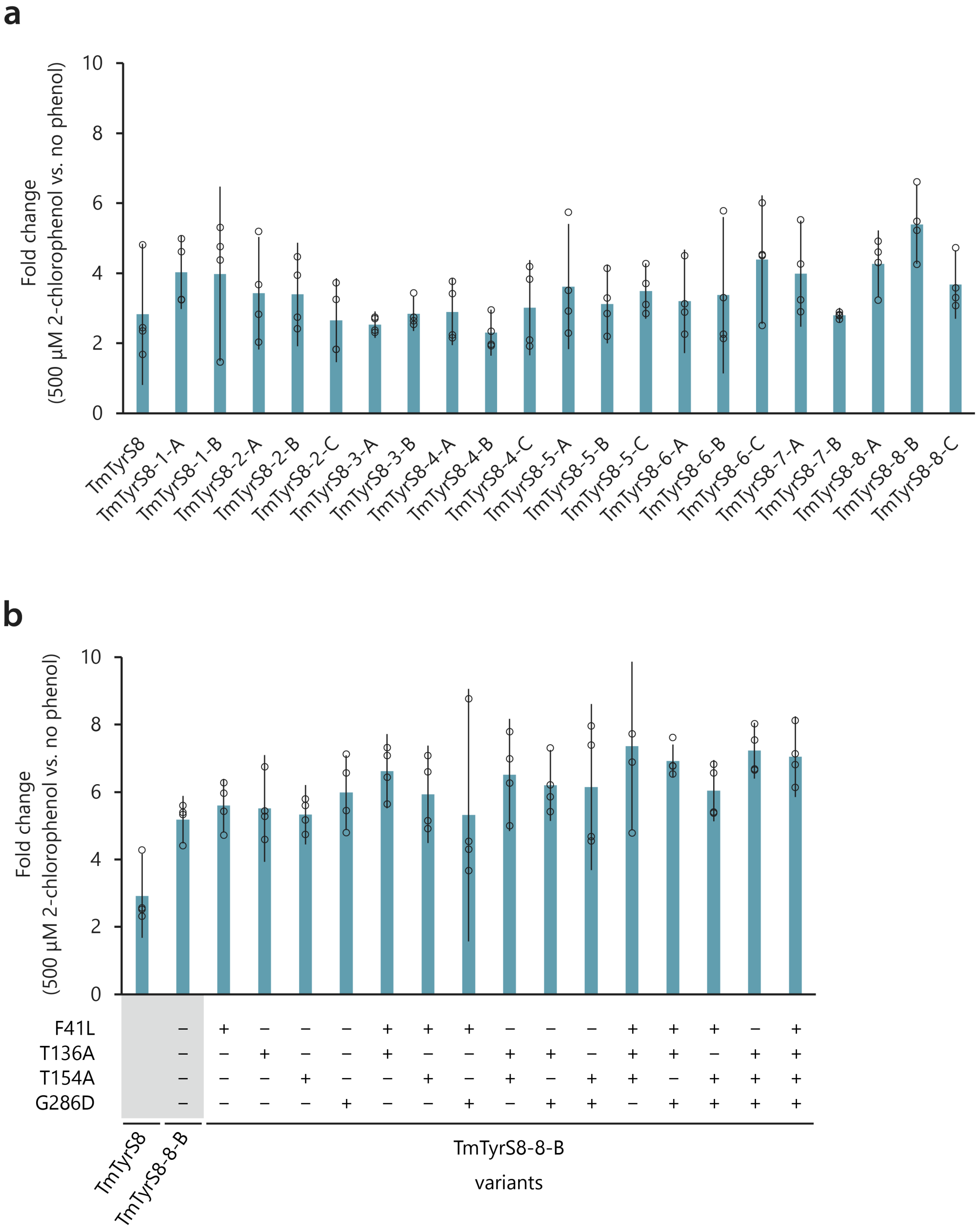


**Supplementary Fig. 3.** Characterization and engineering of *Tm*TyrS8-derived variants in response to 2-chlorophenol. (a) Fold increase in RREs relative to no phenol in yeast cells expressing variants obtained from the *Tm*TyrS8 evolution campaign in the presence of 500 µM 2-chlorophenol. (b) Fold increase in RREs relative to no phenol for engineered variants based on *Tm*TyrS8-8-B, the best performing variant in panel (a). Beneficial mutations were extracted from other variants that exhibited enhanced responses relative to *Tm*TyrS8, yielding substitutions F41L, T136A, T154A, and G286D. Plus and minus symbols in the graph labels indicate the presence or absence of each mutation in the corresponding variant. *Tm*TyrS8-8-B carrying F41L, T136A, and T154A was designated as *Tm*TyrS9. RRE values measured in the presence of 500 µM 2-chlorophenol were normalized to those measured in its absence. Each condition was measured in biological quadruplicates, and the mean ± one standard deviation (error bars) is shown.

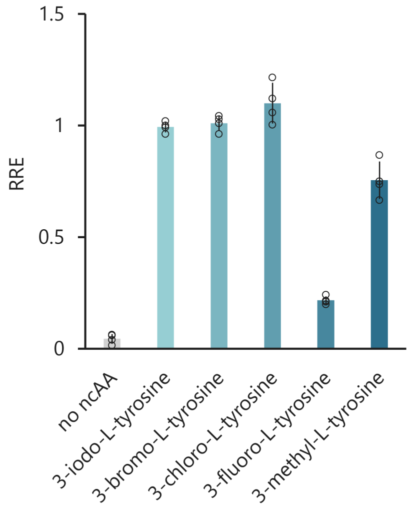


**Supplementary Fig. 4.** ncTyr incorporation by NitroY-F5/3FY-D with individual target ncTyr. RRE value with target 500 μM ncTyr is shown. Each condition was measured in biological quadruplicates, and the mean ± one standard deviation (error bars) is shown.
